# Anthropogenic habitat alteration leads to rapid loss of adaptive variation and restoration potential in wild salmon populations

**DOI:** 10.1101/310714

**Authors:** Tasha Q. Thompson, Renee M. Bellinger, Sean M. O’Rourke, Daniel J. Prince, Alexander E. Stevenson, Antonia T. Rodrigues, Matthew R. Sloat, Camilla F. Speller, Dongya Y. Yang, Virginia L. Butler, Michael A. Banks, Michael R. Miller

## Abstract

Phenotypic variation is critical for the long-term persistence of species and populations. Anthropogenic activities have caused substantial shifts and reductions in phenotypic variation across diverse taxa, but the underlying mechanism (i.e., phenotypic plasticity and/or genetic evolution) and potential to recover previous phenotypic characteristics are unclear. Here we investigate changes in adult migration characteristics of wild salmon populations caused by dam construction and other anthropogenic habitat modifications. Strikingly, we find that dramatic allele frequency change (i.e., genetic evolution) from strong selection at a single locus explains the rapid phenotypic shift observed after recent dam construction. Furthermore, ancient DNA analysis confirms the abundance of a specific allele associated with adult migration type in historical habitat that will soon become accessible through a large restoration (i.e., dam removal) project. However, analysis of contemporary samples suggests the restoration will be challenged by loss of the allele from potential source populations. These results highlight the need to conserve and restore critical adaptive variation before the potential for recovery is lost.

## Introduction

Phenotypic variation buffers species and populations against environmental variability and is important for long-term persistence (1–7). In phenotypically diverse populations, environmental fluctuations that negatively impact one phenotype may have a neutral or positive impact on another (5, 8). This decreases variance in population size across time and reduces vulnerability to extirpation or extinction. Furthermore, phenotypic variation increases the potential for species to persist through long-term environmental changes (e.g., climate change) by serving as the substrate upon which evolution can act. Thus, maintaining intraspecific phenotypic variation is an important component of biodiversity conservation.

Anthropogenic activities have major effects on phenotypic variation across a broad array of species and traits, often producing substantial phenotypic shifts and reductions in overall variation (5, 6, 9–12). Despite the recognized importance of intraspecific variation, the urgency of addressing human-driven phenotypic change through conservation policy and action is unclear because the ability of populations and/or species to recover previous characteristics (e.g., variation) is not well understood (5, 13, 14). If previous variation can quickly reemerge, human-induced phenotypic change may have limited impact on long-term persistence and evolutionary potential. On the other hand, if human actions cause more permanent changes and reductions in variation, immediate steps to reduce impacts on intraspecific phenotypic variation may be warranted.

The mechanisms that underlie human-induced phenotypic change (i.e., phenotypic plasticity and/or genetic evolution) will influence the potential for recovery. For example, if phenotypic changes are due to plasticity (i.e., the ability of the same genotype to produce different phenotypes when exposed to different environments), previous characteristics may rapidly reemerge if environmental conditions change (e.g., habitat is restored) (15, 16). On the other hand, phenotypic change due to genetic evolution (i.e., changes in allele and genotype frequencies across generations) may severely impact the ability to recover previous characteristics (5, 12, 17). In the case of genetic evolution, the ability to recover previous phenotypic characteristics will depend on factors such as the genetic architecture of the affected trait (18). Unfortunately, understanding the genetic basis of phenotypic variation, and thus the potential consequences of human-drive phenotypic change, can be challenging because the genes that influence specific traits in natural populations are usually unknown (19, 20).

The adult migration characteristics of Chinook salmon (*Oncorhynchus tshawytscha*) is a clear example of adaptive phenotypic variation that has been impacted by anthropogenic activities (11, 21, 22). Across the southern part of their coastal (i.e., non-interior) range in North America, Chinook display two primary phenotypes in the characteristics of their spawning migration (23). Premature migrating Chinook enter freshwater from the ocean in a sexually immature state during the spring, migrate high into watersheds to near their spawning grounds, and hold over the summer in a fasted state while their gonads develop before spawning in the fall. Mature migrating Chinook enter freshwater in a sexually mature state in the fall and migrate directly to their spawning grounds to spawn immediately (23). The premature and mature migrating phenotypes are commonly referred to as “spring-run” and “fall-run”, respectively, which will be the nomenclature used here. Although complex phenotypic differences exist between spring-run and fall-run Chinook, freshwater entry date can serve as a good proxy when more extensive measurements (e.g., gamete maturation state and body fat content at freshwater entry, time between freshwater entry and spawning, etc.) are not available (23, 24). The spatial and temporal differences between the two migration types facilitate use of heterogeneous habitats and provide resilience against environmental variability (2, 23, 25).

Many rivers historically hosted large numbers of both phenotypes (26, 27). However, because they rely on clean, cold water throughout hot summer months, spring-run Chinook are more vulnerable than fall-run Chinook to anthropogenic activities that affect river conditions such as logging, mining, dam construction, and water diversion (11, 13, 23, 26, 28). Consequently, in locations where both phenotypes existed historically, the spring-run phenotype has either dramatically declined in relative frequency or disappeared completely since the arrival of Europeans (21, 29). Despite the recognized cultural, ecological, and economic importance of spring-run Chinook (30) as well as their significant contribution to resilience and evolutionary potential, the widespread declines and extirpations of spring-run Chinook have been met with limited conservation concern because previous research suggested that the spring-run phenotype could rapidly reemerge from fall-run populations if habitat conditions improved (13, 31). Here we investigate the mechanism underlying the dramatic decline of the spring-run phenotype and its future recovery potential.

## Results

### Rapid genetic change from strong selection at a single locus explains phenotypic shift in Rogue Chinook

As one of the few remaining locations with a significant number of wild spring-run Chinook (32), the Rogue River in Oregon (Figure 1A) presents a prime opportunity to examine the mechanism behind anthropogenically-induced changes in Chinook migration characteristics. Prior to construction of Lost Creek Dam (LCD) in 1977, Chinook entered the upper basin (i.e., crossed Gold Ray Fish Counting Station [GRS]) almost exclusively in the spring. After dam construction, the Chinook population experienced a phenotypic shift that, by the 2000s, had resulted in a striking increase in the number of individuals entering the upper basin in summer and fall, and a corresponding decrease in the number entering in the spring (Figure 1B; Table S1) (22). This shift occurred despite the majority of Chinook spawning habitat existing below the dam site (22). Because the dam altered downstream temperature and flow regimes (Figure S1), this shift may have resulted from phenotypic plasticity, where post-dam environmental conditions cue fish to migrate later. Alternatively or in addition, the phenotypic shift may have resulted from rapid genetic evolution due to selection caused by post-dam conditions.

**Figure 1.**
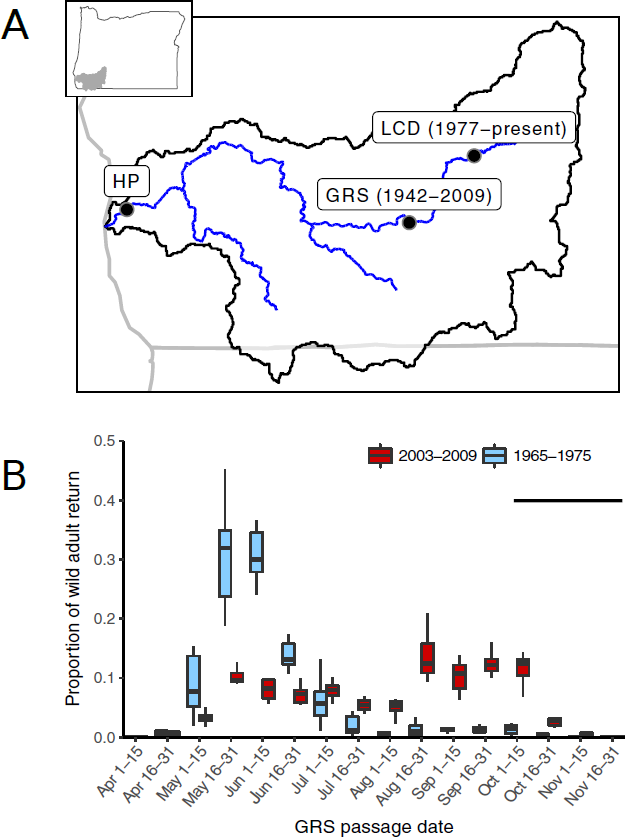
Phenotypic change in Rogue River Chinook. (A) Map of Rogue River; HP: Huntley Park; GRS: Gold Ray Fish Counting Station; LCD: Lost Creek Dam; dates indicate presence of features. (B) Bimonthly proportion of annual wild adult Chinook return across GRS before (1965-1975, 1968 was excluded due to incomplete data) and after (2003-2009, counts prior to 2003 included hatchery fish and GRS was removed in 2010) LCD construction; horizontal bar depicts Chinook spawn timing.

To begin investigating the shift in Rogue Chinook migration characteristics, we analyzed 269 fish that crossed GRS during three approximately week-long intervals in late May (n=88), early August (n=89), and early October (n=92). Each fish was genotyped at the *GREB1L* locus, which was previously found to be associated with migration type (i.e., spring-run or fall-run) across a wide array of Chinook populations (30), using a newly developed marker (see Materials and Methods; Table S2, S5). Strikingly, the three groups had dramatically different genotype frequencies (Figure 2A). All but one late May fish were homozygous for the allele associated with the spring-run phenotype, with the single heterozygote passing GRS on the last day of that collection period (Figure 2B; Table S3). The majority of early August fish were heterozygous. Interestingly, although the early October group was overwhelming homozygous for the fall-run allele, a few individuals were heterozygous or even homozygous for the spring-run allele (Figure 2A). GRS is located approximately 200 km from the river mouth and thus the heterozygous and homozygous spring-run fish that passed GRS in early October may have entered freshwater earlier but held below GRS for an extended period before passage. We conclude that there is a strong association between the *GREB1L* genotype and GRS passage date in Rogue Chinook and that heterozygotes have an intermediate migration phenotype.

**Figure 2.**
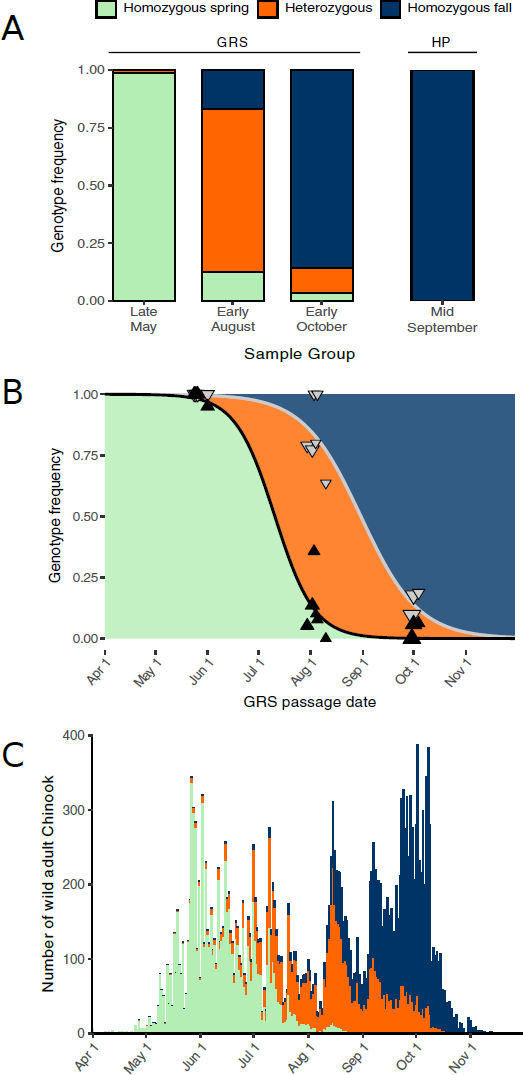
Genetic basis of adult migration phenotype in Rogue River Chinook. (A) Stacked bar graph representing observed *GREB1L* genotype frequencies in GRS and HP sample groups. (B) Scatterplot representing observed *GREB1L* genotype frequencies in GRS samples across 13 collection days; triangles represent homozygous spring-run (black) and homozygous spring-run plus heterozygous (grey) genotype frequencies; triangle size is proportional to the number of fish analyzed each day (min 10, max 42). For fish that pass GRS during a specific time interval (e.g., a single day), the area below the black line represents the expected frequency of the homozygous spring-run genotype, the area between the lines represents heterozygotes, and above the gray line represents the homozygous fall- run genotype. (C) Stacked bar graph representing number of wild adult Chinook passing GRS in 2004; colors represent estimated proportion of each *GREB1L* genotype.

To further investigate the association between *GREB1L* and the migration characteristics of Rogue Chinook, we genotyped 38 fish collected in mid-September at Huntley Park (HP; Figure 1A). HP is located on the mainstem Rogue approximately 13 km from the river mouth so, unlike GRS samples, HP fish are unlikely to have been in freshwater for an extended period prior to collection. Strikingly, all HP samples were homozygous for the fall-run allele (Figure 2A), a significantly lower homozygous spring-run/heterozygous genotype frequency than GRS early October samples (p-value=0.003; binomial distribution). This suggests that heterozygous and homozygous spring-run fish from GRS in early October likely entered freshwater earlier in the year but held for an extended period below GRS before crossing. We conclude that genotype at the *GREB1L* locus is a better predictor of migration type (spring-run, fall-run, or intermediate) than passage date at GRS.

We next estimated the total number of fish of each genotype that passed GRS by extrapolating the genotype frequencies across the entire run year. Briefly, we fit the genotype frequencies with sigmoidal curves to estimate the probability that a fish ascending GRS on any specific day would be each of the three possible genotypes (Figure 2B). We then multiplied the observed number of individuals passing on each day by the genotype probabilities for the same day (Figure 2C; Table S1). Lastly, we performed bootstrap resampling of the daily genotype data to determine 95% confidence intervals for this and subsequent analyses. The analysis suggested that, of the 24,332 individuals that passed GRS in 2004 (Table S1), 8,561 (7,825-9,527) were homozygous for the spring-run allele, 6,636 (5,077-7,798) were heterozygous, and 9,135 (8,124-10,253) were homozygous fall-run. These abundance estimates correspond to homozygous spring-run, heterozygous, and homozygous fall-run genotype frequencies of 0.352 (0.322-0.392), 0.273 (0.209-0.320), and 0.375 (0.334-0.421), respectively, as well as a spring-run allele frequency of 0.488 (0.457-0.518) and a fall-run allele frequency of 0.512 (0.482-0.543). Notably, the estimated homozygous spring-run migration date distribution was strikingly similar to the empirical migration date distribution prior to LCD construction (Figure 1B, 2C), suggesting the pre-dam population was predominantly homozygous spring-run and the migration time of this genotype has not changed since dam construction. This was further supported by an analysis of 36 pre-dam samples collected near the historical late-May/early-June GRS migration peak (Figure 1B), all of which were homozygous for the spring-run allele (see Materials and Methods; Table S3). We conclude that the phenotypic shift after dam construction is explained by rapid allele and genotype frequency shifts at the *GREB1L* locus.

To explore selection regimes that could produce this genetic change in such a short time frame (approximately 7 generations), we estimated the spring-run allele frequency prior to LCD and the selection coefficients required to reach the observed 2004 allele frequency under a simple model assuming the spring-run allele was either recessive, dominant, or codominant with respect to fitness (18). Under the recessive scenario, heterozygous and homozygous fall-run genotypes have equal fitness (selection coefficients: s_FF_=s_SF_=0, 0≤s_SS_≤1). Under the dominant scenario, heterozygous and homozygous spring-run genotypes have equal fitness (s_FF_=0, 0≤s_SF_=s_SS_≤1). Under the codominant scenario, heterozygotes have an intermediate fitness (sFF=0, sSF=½sSS, 0≤sSS≤1). Applying the genotype probability distribution (Figure 2B) to the pre-dam fish counts (Figure 1B) suggested a pre-dam spring-run allele frequency of 0.895 (0.873-0.919), which the pre-dam sample analysis supports as a reasonable estimate (see Materials and Methods; Table S3). Next, the modeling estimated selection coefficients for the homozygous spring-run genotype (sSS) of 0.367 (0.348-0.391), 0.646 (0.594-0.712), and 0.447 (0.424-0.480) under the recessive, dominant, and codominant scenarios, respectively. Furthermore, assuming the same environmental conditions (i.e., selection coefficients) continue into the future, the modeling predicted the spring-run allele frequency in 2100 would be 0.106 (0.099-0.112), 3.24×10-11 (2.44×10-13 – 7.96×10-10), and 0.002 (0.001-0.003) under the recessive, dominant, and codominant scenarios, respectively (Figure 3). Thus, our modeling demonstrates that selection strong enough to explain these rapid phenotypic and genotypic shifts could lead to loss of the spring-run allele in a relatively short time. We conclude that, under continual selection against the spring-run phenotype, the spring-run allele cannot be expected to persist unless recessive with respect to fitness.

**Figure 3.**
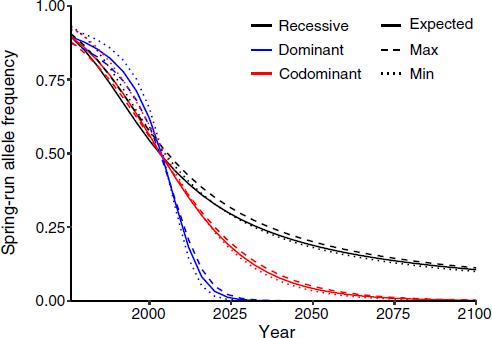
Selection modeling in Rogue Chinook. Line graph representing the spring-run allele frequency over time under recessive, dominant, and codominant scenarios. Estimated spring-run allele frequencies in 1976 (one year prior to LCD construction) and 2004 were used to determine selection coefficients for each scenario (recessive: sFF=sSF=0, sSS=0.367; dominant: sFF=0, sSF=sSS=0.646; codominant: sFF=0, sSF=½(sSS), sSS=0.447). The modeling assumes random mating and no genetic drift.

### Ancient and contemporary Klamath Chinook reveal hindered spring-run restoration potential

The Klamath River in Northern California and Southern Oregon (Figure 4) historically hosted hundreds of thousands of adult spring-run Chinook annually, with the spring-run phenotype possibly exceeding the fall-run phenotype in frequency (27). While the fall-run phenotype remains relatively abundant, dam construction and habitat degradation beginning in the late 1800‘s led to severe declines in the spring-run phenotype, with virtually complete loss of wild spring-run Chinook in the mainstem and tributaries except the Salmon River (21, 33). In the last decade, annual returns of wild fall-run Chinook in the Klamath have numbered in the tens to hundreds of thousands (34), while Salmon River spring-run Chinook have ranged from approximately 200 to 1,600 individuals (35) and are expected to be extirpated within 50 years (21). In 2021, the largest-scale dam removal project in history is scheduled to remove four dams in the upper basin (36) and reopen hundreds of miles of historical Chinook habitat inaccessible since 1912 (37) (Figure 4). This dam removal provides an opportunity unprecedented in scale to restore extirpated populations, including spring-run Chinook (38). However, while historical documentation suggests the presence of early-migrating Chinook in the upper Klamath (37), the extent to which above dam populations relied on the same spring-run allele as the Rogue (see above) and other contemporary Chinook populations (30) (see Materials and Methods) is unknown. Furthermore, since most contemporary Klamath populations have lost the spring-run phenotype, it is unclear which, if any, have maintained the spring-run allele and therefore could serve as a source population for restoration of spring-run Chinook in the upper basin.

**Figure 4.**
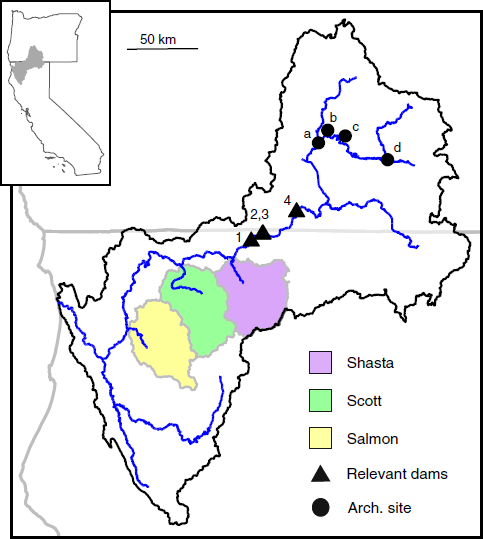
Map of Klamath Basin. Klamath Dams scheduled for removal in 2021: 1) Iron Gate; 2) Copco 1; 3) Copco 2; 4) J.C. Boyle. Archaeological site locations of ancient samples: a) Williamson River Bridge; b) Bezuksewas Village; c) Kawumkan Springs Midden; d) Beatty Curve.

To investigate the genetic composition of historical upper Klamath Chinook, we genotyped nine Chinook samples collected from four archaeological sites in the upper basin known to be historically important fishing places for Klamath peoples (39) (Figure 4). The samples ranged in age from post-European contact to approximately 5,000 years old and, based on the presence of all body parts in the archaeological sites, were likely caught locally as opposed to being acquired through trade (39–41) (Table 1). Strikingly, three of the locations had only homozygous spring-run samples, while the remaining location had only homozygous fall-run samples (Table 1). The spring-run sample locations are known to have been occupied by humans in the spring or throughout the year and are also near major cold-water input sources (suitable over-summering habitat for spring-run Chinook (42)) whereas the fall-run samples came from a location with a documented historical fall fishery (43). We conclude that the upper basin harbored the same allelic variants as contemporary populations, and these spring-run alleles are expected to be necessary for restoration of the spring-run phenotype in the upper basin (see above) (30).

**Table 1.**
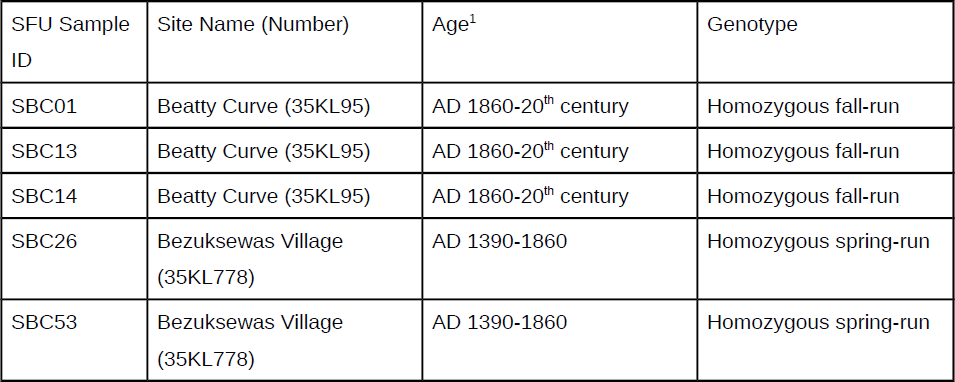
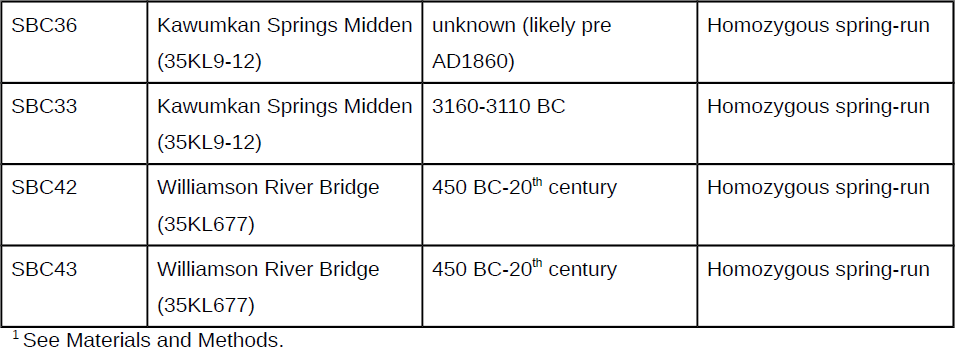
Ancient upper Klamath Chinook sample information and genotyping results, listing Simon Fraser University (SFU) sample identification number and Oregon state site numbers

To test if spring-run alleles are being maintained in lower (i.e., below dam) Klamath populations that have lost the spring-run phenotype, we genotyped juvenile Chinook collected from the Shasta River throughout the juvenile out-migration season in 2008-2012 (Table S4) (44). The Shasta, where spring-run Chinook were last observed in the 1930‘s (27), is a major Klamath tributary that shares many environmental characteristics with the habitat above the dams (e.g., spring water source, dry climate, etc.) (45). Thus, Shasta Chinook may contain additional adaptive variation suitable for the upper Klamath, which makes them an attractive restoration stock candidate (46). Strikingly, out of the 440 successfully genotyped individuals, only two were heterozygous and all others were homozygous for the fall-run allele, corresponding to a spring-run allele frequency of 0.002 (binomial distribution 95% CI: 3×10^−4^ – 0.008; Table 2). This is at least an order of magnitude below the expected frequency if the spring-run allele was recessive with respect to fitness (see Discussion; e.g., Figure 3) and, interestingly, very similar to the codominant scenario in our Rogue Chinook modeling (0.002 vs. 0.002; Figure 3) after a similar period of selection against the spring-run phenotype (late 1800s-early 2000s vs. 1977-2100). Given the recent annual adult returns to the Shasta River and N_e_/N ratios in Chinook (47), such frequencies suggest the spring-run allele is highly vulnerable to complete loss, even without continued selection against heterozygotes (see Discussion). We conclude that the spring-run allele is not being maintained in Shasta Chinook.

**Table 2.** Klamath Chinook smolt information and genotyping results

| River | Date last<br>spring-run<br>Chinook<br>observed | Number | Year(s) | Homozygous<br>spring-run | Heter<br>ozygo<br>us | Homozygous<br>fall-run | Spring-run<br>allele frequency |
| --- | --- | --- | --- | --- | --- | --- | --- |
| Shasta | 1930s† | 440 | 2008-2012 | 0 | 2 | 438 | 0.002 (3e-4 -<br>0.008)* |
| Scott | 1970s | 432 | 2007-2013 | 0 | 2 | 430 | 0.002 (3e-4 -<br>0.008)* |
| Salmon | present | 116 | 2017 | 14 | 19 | 83 | 0.20 |

\*95% CI calculated using binomial probability distribution; †spring-run Chinook were still observed just upstream of the Shasta River mouth at Iron Gate Dam into the 1970s.

To test if the spring-run allele is being lost from locations with disparate environmental conditions, we genotyped Chinook juveniles collected over a similar time range in the Scott River (Figure 4), a Klamath tributary that exhibits a hydrologic regime driven by surface water which is typical of the lower Klamath basin (45). The spring-run phenotype was last observed in the Scott River in the 1970‘s (27). We also genotyped 116 juveniles from the Salmon River (see above; Figure 4) as a positive control. Out of 432 successfully-genotyped Scott samples, we found only two heterozygotes (spring-run allele frequency: 0.002; binomial distribution 95% CI: 3×10^−4^ – 0.008), whereas the Salmon River samples had an overall spring-run allele frequency of 0.20 (Table 2), corresponding well with spring-run Chinook frequency estimates based on annual dive and carcass surveys in the Salmon River (35, 48). We conclude that spring-run alleles are not being maintained in the Scott River and that diverse environments are susceptible to rapid loss of the spring-run allele upon extirpation of the spring-run phenotype.

## Discussion

Complex phenotypic variation (e.g., life history variation) facilitates species resilience in heterogeneous or variable environments (2, 5). The genetic architecture of complex variation, though usually unknown, is typically assumed to also be complex (i.e., polygenic) (49). A recent study identified a single locus (*GREB1L*) associated with migration type in Chinook as well as the closely related species steelhead (*Oncorhynchus mykiss*) (30). However, the relatively low marker resolution and poor phenotypic information in the Chinook analysis obscured the strength of association and phenotype of heterozygotes. Our analysis of samples with more detailed phenotypic information (i.e., specific migration dates at GRS and Huntley Park [see Results; Table S3] as well as the lower South Fork Trinity [see Materials and Methods; Table S5]) using a new marker identified through a high-resolution, multi-population analysis of *GREB1L* (see Materials and Methods; Table S2, S5) suggests that 1) the association of migration type with variation at *GREB1L* is extremely robust and 2) heterozygotes have an intermediate migration phenotype (Figure 2A). Therefore, while phenotypic variation within each genotype (e.g., precise freshwater entry and spawning dates) is yet to be explained, migration type (i.e., premature/spring-run or mature/fall-run) appears to have a strikingly simple genetic architecture. Furthermore, the association of a single haplotype with the spring-run phenotype in diverse locations (Table S5) supports a previous conclusion that spring-run alleles arose from a single evolutionary event and cannot be expected to readily re-evolve (30, 50). Thus, simple modes of inheritance and rare allelic evolutionary events can underpin complex phenotypic variation.

Selection results from the balance between benefits and costs of specific phenotypes (51), and anthropogenic habitat alteration can potentially disrupt this balance (9, 12, 52, 53). The large and rapid decline in the Rogue spring-run phenotype and allele frequency suggests strong selection against spring-run Chinook after LCD construction. Furthermore, our modeling demonstrates how such selection, if sustained, could rapidly result in complete loss of the spring-run allele. A main benefit of the spring-run phenotype is thought to be access to exclusive temporal and/or spatial habitat, while a major cost is reduced gametic investment (e.g., smaller egg size) because energy must be dedicated to maintenance and maturation while fasting in freshwater (23, 54). River flow regimes can be a major driver of life history evolution in aquatic systems (12, 53), and LCD altered temperature and flow in a way that may allow fall-run Chinook access to spawning habitat that was previously exclusive to spring-run Chinook (22). An analysis of carcass samples from the Rogue revealed substantial spatial and temporal overlap in spawning distributions of all three genotypes (Figure S2; Table S3), supporting the hypothesis that anthropogenically-induced habitat alterations have reduced the historical benefit of the spring-run phenotype, contributing to its decline. Regardless of exact mechanisms, our results provide a clear example where anthropogenic factors induced rapid phenotypic change through genetic evolution as opposed to phenotypic plasticity.

Population genetics theory and our selection modeling predicts that, for simply-inherited traits, alleles promoting negatively-selected phenotypes will be eliminated from a population unless they are masked in the heterozygous state (i.e., recessive with respect to fitness) (18). The intermediate migration phenotype of heterozygotes, in combination with typical lower river conditions at intermediate times (i.e., conditions inhospitable to salmonids), suggests their fitness will be at least somewhat lower than fall-run Chinook in most locations (55). Therefore, where the spring-run phenotype is lost, spring-run alleles should not be expected to be maintained in the heterozygous state. This prediction is empirically supported by our results from the Shasta and Scott rivers where, based on adult run size estimates during the years our samples were spawned, the observed spring-run allele frequency (0.002) would correspond to an average of approximately 20 heterozygous adults per year in each river (56, 57). Given that adult Chinook have highly variable reproductive success (47) and our samples were collected prior to the recent extreme drought in California (21), such a low observed frequency makes it plausible spring-run alleles have already been completely lost owing to continued selection and/or genetic drift. Notably, while habitat alterations extirpated the spring-run phenotype from the Shasta and Scott, the total Chinook census sizes (i.e., adults of any migration type) of both rivers are considered robust (56, 57). Thus, both theory and empirical evidence suggest heterozygotes are not a sustainable reservoir for spring-run alleles, and human factors can eliminate important adaptive variation regardless of total population size.

Adaptive variation is likely important to the success of species restoration efforts (46, 58). The planned removal of Klamath dams provides an opportunity to restore Chinook to historical habitat that is unprecedented in scale. Our analysis of ancient samples corroborates historical documentation suggesting both migration types existed above the dams (37). While abundant Klamath fall-run Chinook are likely to naturally recolonize the upper basin, our results suggest the spring-run allele frequency may be too low for natural recolonization of spring-run Chinook. Furthermore, human-facilitated restoration may be challenged by limited options for appropriate source populations (i.e., populations that contain the spring-run allele). The Shasta River‘s environmental similarities with the upper basin (45) would have made it an attractive candidate if spring-run alleles were being maintained (46, 59). Salmon River spring-run Chinook are severely depressed in number (21, 35) and may lack other adaptive variation important for the upper basin due to the major environmental differences between the locations (45, 59). Spring-run alleles exist in a within-basin hatchery population (i.e., Trinity River Hatchery), but hatchery salmonids are partially domesticated, have reduced reproductive success in the wild, and negatively impact wild populations (60–62). Introducing an out-of-basin wild stock (i.e., Rogue spring-run Chinook) is an option but may also be challenged by adaptive incompatibilities (59). Given that wild spring-run Chinook are expected to disappear from the lower Klamath within 50 years and are declining across their range (21), the current challenges of restoring spring-run Chinook upon Klamath dam removal should be considered a preview of even greater challenges that will be faced in future spring-run Chinook restoration projects if the spring-run phenotype continues to decline. Thus, the decline and loss of adaptive variation due to anthropogenic habitat alterations hinders the ability to restore wild populations.

Although this study provides important insights into the genetics and conservation of spring-run Chinook, additional information would be useful to further inform conservation and restoration actions. Future work should broadly characterize the distribution of spring-run alleles, especially in populations that appear to lack the spring-run phenotype, in order to identify if and where the genetic potential for premature migration still exists (e.g., in heterozygotes). Care should be taken so that sampling design accounts for phenotypic differences between genotypes to prevent biased frequency estimates. External factors that may influence allele frequencies (e.g., introgression with a hatchery stock) should also be considered. Ongoing monitoring of allele frequencies will likely also be necessary, as spring-run alleles may be present but in decline. Importantly, a better understanding of the ecology (i.e., spawning and rearing locations), phenotype (i.e., range of river entry and spawning dates, fecundity, etc.), and fitness (i.e., relative reproductive success) of each genotype would be useful for understanding selection mechanisms and targeting conservation strategies. Lastly, although the genetic marker used here is currently the best available to distinguish between migration types (Table S5), continued marker development (e.g., identification of the causative polymorphism[s]) would reduce the potential for misclassification of migration type due to factors such as rare recombination events.

The combination of results from this study provides important insights into the mechanisms and consequences of phenotypic change induced by anthropogenic habitat alteration. First, our results demonstrate that complex phenotypic variation can have a simple genetic architecture and that anthropogenically-induced phenotypic change can be caused by rapid genetic evolution from strong selection at individual loci. Furthermore, our results (both modeled and empirical) demonstrate this situation can lead to the rapid loss of important adaptive alleles, including from populations that are healthy from a total population size perspective. In cases where adaptive alleles are the product of mutational events that are very rare from an evolutionary perspective (such as the spring-run allele in Chinook (30)), their loss will create a major challenge for future restoration as well as limit resilience and evolutionary potential. Taken together, our results highlight the need to conserve and restore critical adaptive variation before the potential for recovery is lost.

## Materials and Methods

### GREB1L marker discovery

Previous research identified a significant association between variation in the *GREB1L* region and adult migration type (i.e., premature or mature) in both Chinook and steelhead (*Oncorhynchus mykiss*) (30, 63). Although the strongest associated SNP in Chinook (position 569200 on scaffold79929e) had a large allele frequency difference between premature and mature migrating populations in several locations (30), this association was notably weaker than observed in steelhead. We reasoned the weaker association could have resulted from technical reasons (e.g., lower SNP resolution of the Chinook analysis) as opposed to biological reasons (e.g., smaller influence of the *GREB1L* locus in Chinook compared to steelhead).

We therefore used capture baits to isolate and sequence the *GREB1L* region in 64 Chinook samples (across 8 locations in California, Oregon, and Washington; Table S5) from the previous association study (30) for additional SNP identification and association testing. The two most strongly associated SNPs identified by this process (positions 640165 and 670329 on scaffold79929e) were approximately 30 kb apart just upstream of *GREB1L* and revealed much stronger associations than the most strongly-associated SNPs from the previous study (30) (Table S5). These results confirm that the relatively weak association between *GREB1L* and migration type previously observed in Chinook (compared to steelhead) (30) was due to lower SNP resolution as opposed to a smaller influence on phenotype.

### SNP assay design and validation

We designed TaqMan-based genotyping assays for the two newly discovered SNPs to facilitate rapid and inexpensive genotyping of the *GREB1L* locus across large numbers of samples. Approximately 300 bp of Chinook sequence surrounding each SNP (Table S2) was submitted to the Custom TaqMan Assay Design Tool (Applied Biosystems) to generate primer and probe sequences for each SNP. Additional polymorphic sites in the surrounding sequence identified in the capture sequencing were masked to avoid primer or probe design across these sites. Assays were run using 5 µl 2X TaqMan Genotyping Master Mix, 0.5 µl 20X genotyping assay (final concentrations of 900 nM [primers] and 200 nM [MGB probes]), 2.5 µl DNA-grade water, and 2 µl sample DNA for each reaction. Reporter dyes were Vic and Fam. Each 96-well SNP assay plate also contained one positive control for each genotype (taken from samples used in capture sequencing) and two negatives controls substituting water or low TE (0.1 mM EDTA, 10 mM tris pH 8.0) for DNA. No negative controls ever amplified. Each SNP assay was run separately (not multiplexed) for each sample. The assays were run on either a Chromo4 or QuantStudio-3 Real Time PCR machine for 10 minutes at 95°C followed by 40 cycles of 15 seconds at 95°C and 1 minute at 58-59°C (snp640165) or 62-64°C (snp670329).

SNP assays were validated with the samples used for capture sequencing. All results were consistent with sequencing-based genotype calls (Table S5). Our genotyping results from GRS and Huntley Park (Figure 2A; Table S3) serve as further validation of the assays in the Rogue River. For additional validation in the Klamath, we genotyped 62 samples from Chinook with known migration dates through a weir on the lower South Fork Trinity River (Table S5). All South Fork Trinity samples phenotyped as spring-run (i.e., weir passages dates between mid-May and end of July) were homozygous for the spring-run allele except for a single heterozygote collected on July 31. All samples phenotyped as fall-run (i.e., weir passages dates between mid-October and mid-November) were homozygous for the fall-run allele (Table S5).

### Contemporary sample collection and DNA extraction

Rogue GRS samples were obtained from wild Chinook salmon, defined as lacking an adipose fin clip, that returned to spawn in the Rogue River during 2004. Fish were trapped by Oregon Department of Fish and Wildlife (ODFW) personnel at a fish-count station (GRS) located at Gold Ray Dam (erected in 1941). Tissue was sampled from the operculum of each fish and placed in 100% ethanol for storage and subsequent DNA extraction using Qiagen DNeasy kits following manufacturers protocols. Following sampling, fish were released unharmed upstream of the dam barrier. Approximately 300 samples were evenly obtained across three temporal sampling windows (May 24 to June 1; July 30 to August 10; and September 30 to October 4) that targeted spring, intermediate, and fall runs.

Rogue Huntley Park samples were collected from wild Chinook caught in beach seines near Huntley Park in September 2014 (Table S3). Rogue pre-LCD samples were collected in the lower river in during May of 1975 and 1976 (Table S3) and stored in the ODFW scale archive. Rogue carcass samples were collected during ODFW spawning surveys of the upper Rogue in 2014 (Table S3). Juvenile Chinook from the Salmon, Shasta, and Scott Rivers in the Klamath Basin were caught in screwtraps during smolt outmigration across several years (Table S4) (44). South Fork Trinity samples were collected from live adult Chinook during passage through Sandy Bar weir, except for three samples that were collected at Forest Glen (Table S5). Fin clip (Huntley Park, Rogue carcass, and Salmon) or scale (Rogue pre-LCD, Shasta, and Scott) samples were collected, dried on filter paper, and stored at room temperature. DNA was extracted using a magnetic bead-based protocol (64) and stored at −20°C.

### Archaeological sample collection and DNA extraction

The archaeological samples were recovered from archaeological excavation projects led by research teams from the University of Oregon Museum of Natural and Cultural History (Eugene, OR) between the late 1940s and the late 2000s (39, 65). The four sites represent fishing camps or year-round villages occupied by ancestral people to the Klamath Tribes of Oregon (Table 1, S4). Three sites are located on the Sprague River: Kawumkan Springs Midden (66), Beatty Curve (65), and Bezuksewas Village (67). A fourth, Williamson River Bridge (68), is located near the confluence of the Williamson and Sprague Rivers (Figure 4). The sites range in age from 7500 years ago to the early 20^th^ century (39). Because of severe stratigraphic disturbance by burrowing rodents, the materials can typically only be assigned to very broad time periods (Table 1, S4). Deposits were assigned to A.D. 1860 or later based on presence of artifacts of Euro-American origin, as A.D. 1860 marks the establishment of Fort Klamath and time of sustained Euro-American contact in the upper Klamath Basin. Klamath people continued to fish and occupy the Beatty Curve and Williamson River Bridge site locations into the 20^th^ century, so the end date is uncertain. All other ages were based on multiple radiocarbon samples (39), calibrated using OxCal v4.2 (69).

Previous projects (39) assigned the fish remains to the finest taxon possible using modern reference skeletons from known species. To obtain species-level identification, a sample of salmonid remains was sent to the dedicated Ancient DNA Laboratory in the Department of Archaeology at Simon Fraser University, Burnaby, Canada. Twelve vertebra samples (9 Chinook and 3 steelhead as controls) were included in this study (Table S4). Samples were chemically decontaminated through submersion in commercial bleach (4-6% sodium hypochlorite) for 10 minutes, rinsed twice with ultra- pure water, and UV irradiated for 30 minutes each on two sides. Bones were crushed into powder and incubated overnight in a lysis buffer (0.5 M EDTA pH 8.0; 0.25% SDS; 0.5 mg/mL proteinase K) in a rotating hybridization oven at 50°C. Samples were then centrifuged and 2.5-3.0 mL of supernatant from each sample was concentrated to <100 μL using Amicon Ultra-4 Centrifugal Filter Devices (10 KD, 4mL, Millipore). Concentrated extracts were purified using QIAquick spin columns based on previously developed methods (70, 71). 100 μL of DNA from each sample was eluted from QIAquick columns for PCR amplifications.

Species identification was accomplished by targeting salmonid mitochondrial d-loop (249bp) and cytochrome b (cytb) (168bp) fragments as previously described (72). Successfully amplified products were sequenced at Eurofins MWG Operon Ltd. using forward and/or reverse primers. The resulting sequences were compared to Genbank reference sequences through the BLAST application to determine their closest match, and species identifications were confirmed through multiple alignments of the ancient sequences and published salmonid reference sequences conducted using ClustalW (73) through BioEdit (74), as well as the construction of neighbor-joining phylogenetic trees using Kimura‘s 2-parameter model in the Mega 6.0 software program (75). Nine of the 12 samples were identified as Chinook (Table S4), the remaining three as steelhead.

### Rogue and contemporary Klamath genotyping

After DNA extraction, samples were genotyped using the assays (snp640165 and snp670329; Table S2) and qPCR protocol described above. All samples were tested at both SNPs, and a genotype call (homozygous spring-run, heterozygous, or homozygous fall-run; Table S3, S4) was made only if both SNPs were successfully genotyped and consistent with each other. The causative polymorphism(s) at the *GREB1L* locus are currently unknown, so requiring successful and consistent calls at both associated SNPs provides greater confidence that the genotype (homozygous spring-run, heterozygous, or homozygous fall-run) was not miscalled due to biological factors such as rare recombination events and is more conservative than using a single SNP. Of the 1304 samples tested from live-caught fish, 1247 (95.6%) successfully genotyped at both SNPs, 31 (2.4%) failed at one SNP, and 26 (2.0%) failed at both SNPs. Of the 96 Rogue River carcass samples tested, 86 (89.6%) successfully genotyped at both SNPs, 2 (2.1%) failed at one SNP, and 8 (8.3%) failed at both SNPs. Of the successful live and carcass samples (1333 total), 1320 (99%) had the same genotype call at both SNPs, indicating near perfect linkage disequilibrium (LD) between the SNPs. The remaining 13 samples (all from the Rogue [2.8% of successfully genotyped Rogue samples] and mostly from the GRS August group) had a homozygous genotype at one SNP and a heterozygous genotype at the other (Table S3). Because we do not know which, if either, SNP is in stronger LD with the causative polymorphism(s), these samples were called as ambiguous (Table S3) and excluded from further analyses.

### Ancient Klamath genotyping

Multiple sealed aliquots of extracted ancient DNA from 12 archaeological samples were shipped from Simon Fraser University to the University of California, Davis on dry ice. Nine samples were from Chinook and the remaining three were from steelhead, which are known to have the same alleles as fall-run Chinook at the two SNPs based on the *O. mykiss* reference genome (76). Genotyping was conducted under blinded conditions with respect to species, location, and age. SNP assays were run using 10 µl 2X TaqMan Genotyping Master Mix, 1 µl 20X genotyping assay (final concentrations of 900 nM[primers] and 200 nM [MGB probes]), 5 µl DNA-grade water, and 4 µl of sample DNA diluted in low TE (either 1:10 or 1:50) for each reaction. The assays were run on a QuantStudio-3 Real Time PCR machine for 10 minutes at 95°C followed by 80 cycles of 15 seconds at 95°C and 1 minute at 58°C (snp640165) or 64°C (snp670329). Fluorescence after each amplification cycle was measured and checked to prevent erroneous calls due to high cycle number. All plates contained positive controls for each genotype diluted at ratios similar to the unknown samples and at least 12 negative controls substituting the low TE used in sample dilutions in place of DNA. No amplification was ever observed in a negative control in either the ancient sample plates or any plates containing contemporary samples. All results were replicated using separately-sealed aliquots on different days. Due to the extremely high LD in contemporary samples and the precious nature of the ancient samples, genotypes were called even if only one SNP was successfully genotyped (Table S4). Requiring both SNPs to be successfully genotyped would have reduced the number of ancient Chinook samples with a migration type call from nine to five (two fall-run and three spring-run; Table S4) but would not have altered our conclusions.

### Curve fitting and selection modeling

Sigmoidal curves were fit to the genotype frequencies measured for each collection day at GRS (Figure 2B; Table S3). The curves were fit using the Nonlinear Least Squares (nls) function in R (77) for a sigmoidal model, optimizing for b and m values: S = 1/(1 + e-b(x-m)). The R command used was: nls(gf~1/(1 + exp(-b * (x-m))), weights=w, start=list(b=(−0.01), m=90)) where gf was either a list of the homozygous spring-run or homozygous spring-run plus heterozygous frequencies (a.k.a. 1 – homozygous fall-run frequency) with each frequency corresponding to a specific sample collection day, x was a list of numeric dates (April 1 was set to day 1) corresponding to each collection day, and w was the number of samples from each day. The resulting equations represent the estimated probability of each genotype on any given day (Figure 2B), and were applied to daily empirical GRS fish counts from 2004 to estimate allele frequencies in 2004.

Pre-LCD allele frequencies were estimated by applying the genotype probability distribution calculated from the 2004 GRS samples (Figure 2B) to the average bi-weekly fish counts (using mean probability across the bi-weekly bin) in the decade prior to LCD construction, and resulted in a pre- LCD spring-run allele frequency estimate of approximately 90% (see Results). This approach was used because a pre-LCD sample set adequate to perform a direct estimate of the pre-LCD allele frequencies (e.g., pre-LCD samples collected at GRS throughout the migration season) was not available. However, this approach assumes that the relationship between *GREB1L* genotype and GRS passage date was not substantially different pre- and post-LCD. If this assumption is inaccurate (e.g., the association of *GREB1L* with GRS passage date was weaker in the pre-LCD environment), the pre- LCD population may have had a spring-run allele frequency significantly lower than 90%.

We investigated this possibility by genotyping 36 pre-LCD adult Chinook sampled in May (mean date May 20) from the lower Rogue (mean river mile 17) at the *GREB1L* locus (Table S3). Based on measured migration rates of Rogue Chinook (22), these fish would likely have passed GRS near or somewhat after the pre-LCD migration peak in late-May/early-June (Figure 1B). Strikingly, all 36 samples were homozygous for the spring-run allele (Table S3). This demonstrates that pre-LCD individuals that passing GRS around the spring migration peak overwhelmingly contained the spring-run allele and, since very few pre-LCD individuals passed GRS later in the year, suggests our pre-LCD spring-run allele frequency is unlikely to be an overestimate. Furthermore, because the curves are fit to genotype frequencies from post-LCD conditions where heterozygotes are likely more frequent, the pre-LCD allele frequency results likely underestimate the true spring-run allele frequency prior to LCD. Thus, the true change in allele frequency after LCD is probably somewhat greater than what is estimated here, and therefore, our estimated allele frequencies and selection coefficients are likely conservative.

The strength of selection against the spring-run phenotype (i.e., the homozygous spring-run selection coefficient [sSS]) was estimated by calculating values of sSS that explain the estimated change in spring-run allele frequencies between pre-LCD and 2004 using the equation p’ = (sSS p2 + sSF p(1-p)/(sSS p2 + sSF 2p(1-p) + sFF (1-p)2) (18) where sxx is the selection coefficient of each genotype, p is the spring-run allele frequency in the current generation, and p’ is the spring-run allele frequency in the next generation. The estimated pre-LCD spring-run allele frequency was used as the starting value of p, and the equation was run recursively using the p’ value from the current run as the next value of p to find values of sSS that resulted in the estimated 2004 spring-run allele frequency after seven generations (assuming 4-year generations). Calculations were conducted under three relative fitness scenarios: recessive (s_SF_ = sFF), dominant (s_SS_ = s_SF_), and codominant (s_SS_ = 2sSF). The homozygous fall-run genotype was always assumed to have the lowest selection coefficient (s_FF_ = 0). This approach assumes Hardy-Weinberg Equilibrium (HWE), which is probably violated because the slightly earlier mean spawning date of spring-run Chinook likely creates some level of assortative mating (Figure S2). Under assortative mating, the overrepresentation of homozygous spring-run individuals could lead to an even more rapid decrease in the spring-run allele frequency because homozygous spring-run experiences the strongest selection in our modeling. Thus, assuming HWE likely produces conservative selection coefficient and future allele frequency estimates.

## Acknowledgments

We thank C. Bean, R. Bowden, J. Bull, B. Chesney, A. Corum, J. Crawford, M. Johnson, D. Jacobson, L. Ketchum, J. Minch, K. O‘Malley, M. Pepping, B. Quinter, S. Richardson, T. Satterthwaite, T. Soto, and P. Tronquet for help with sample acquisition; T. Satterthwaite for GRS sampling design and fish count data; D. Van Dyke and S. Clements for valuable feedback on initial Rogue results; R. Peak for assistance with map visualization; and M. Hereford and R. Waples for valuable comments on an earlier version of the manuscript. We also acknowledge the help and support of the Klamath Tribes, particularly P. Chocktoot, Jr. and L. Dunsmoor, as well as P. Endzweig, T. Connolly, D. Jenkins, and J. Erlandson (Museum of Natural and Cultural History in Eugene, Oregon) for facilitating access to archaeological fish remains. Partial funding for this work was provided by the Gordon and Betty Moore Foundation through a Science Capacity Grant to the Wild Salmon Center. Part of the ancient DNA work was funded by a National Oceanic and Atmospheric Administration contract (AB133F09CQ0039), through the efforts of J. Simondet and I. Lagomarsino with help from J. Hamilton and B. Tinniswood.

---

Figure S1. Change in Rogue River temperature and discharge following construction of Lost Creek Dam as measured at USGS stream gage site 14337600 near McLeod, Oregon. Lines represent differences in 7-day running averages of maximum daily stream temperature (°C) and mean daily discharge (cubic meters per second) between post- (2003-2009) and pre- (1970-1975) dam periods.

Figure S2. Genotyping results from adult Chinook carcasses recovered in the upper Rogue River during surveys in 2014. Sample locations are shown as the middle kilometer of the survey reach (a stretch of river several kilometers in length) where they were recovered. Lost Creek Dam is approximately 50 km above Gold Ray Fish Counting Station (GRS).

